# OligoSeq: Rapid nanopore-sequencing of single-stranded oligonucleotides

**DOI:** 10.1101/2025.06.10.658886

**Authors:** Robert Ramirez-Garcia, Akashaditya Das, Thomas Heinis

**Affiliations:** Department of Computing, Faculty of Engineering, Imperial College London, London, SW7 2RH GB, United Kingdom; London Biofoundry, Translation and Innovation Hub (I-HUB), Imperial College London, London, W12 0BZ GB, United Kingdom

**Keywords:** DNA sequencing, oligonucleotide sequencing, OligoSeq, AmpliSeq, RevSeq, random primers, nanopore technology

## Abstract

Nanopore-based DNA sequencing technology has achieved remarkable success in sequencing increasingly long DNA strands (e.g. over a million nucleotides long) for genomics research and biotechnology applications. However, the same level of progress has not been achieved for DNA oligonucleotides (usually ≤ 300 nucleotides long). Oligonucleotides play a crucial role in genome engineering efforts through oligo library generation and in DNA data storage, where they are used to encode computer information (such as binary code) in DNA libraries. To enable these applications, accurate sequencing of oligonucleotides in a way that allows to assess for sequence variability, quality and length is essential. But sequencing solutions for oligonucleotides — particularly DNA primers for PCR, oligo DNA libraries used for mutagenesis or cDNA libraries used in gene expression analysis — remain inadequate. To address this gap, we develop *OligoSeq*, an innovative approach that integrates two complementary techniques: AmpliSeq (based on PCR) and RevSeq (based on reverse complementation of either sequence-specific or random primers) to facilitate sequencing of single-stranded oligonucleotides using reference sequence anchor matches of more than ≥90 % identity spanning from about 70 % to 10 % with AmpliSeq or RevSeq with random nonamers, respectively, and resolving the final reference sequence based on the most likely candidate from basecall frequencies, regardless of length and double-stranding method. OligoSeq uses nanopore technology and can be used as a reference for other sequencing platforms requiring double-stranded adapters, offering a practical and scalable alternative for standard quality control in single-stranded oligonucleotide synthesis. The use of nanopore technology, compatible with the double-stranding methods showcased, is shown to be the most cost-effective method for resolving original DNA sequences of different length and quality, and to assess its sequence variability, compared to other methods such as Illumina, PacBio or HPLC/MS.

## Introduction

DNA sequencing technology has made significant advances in sequencing long DNA strands for genomics and biotechnological applications. However, similar progress has not been achieved for shorter sequences, such as RNA transcripts or DNA oligonucleotides. Oligonucleotides play a crucial role in synthetic biology, enabling the creation of extensive libraries for genome engineering [1][2] and serving as a medium for encoding binary data in DNA storage [3].

This underscores the importance of accurately sequencing oligonucleotides of different quality and length. However, oligonucleotide sequencing methods, including those for RNA libraries used in gene expression analysis and protein bioproduction optimization [4], remain neither readily accessible nor standardised. Instead, quality control often depends on expensive techniques such as High Performance Liquid Chromatography (HPLC) and Mass Spectrometry (MS) [5][6][7].

This work presents *OligoSeq*, a comprehensive method for the sequencing of single-stranded oligonucleotides, that goes from double-stranding of single-stranded oligos to sample preparation and sequencing. The double-stranding techniques showcased comprise both `AmpliSeq` and `RevSeq` and offer a promising solution for rapid, accessible, and standardised sequencing of single-stranded oligonucleotides using nanopore technology (Figure 1 and Figure 2). Moreover, it is adaptable to all other sequencing platforms that require flanking adaptors for the analysis of double-stranded nucleic acids. Many widely available DNA sequencing technologies require double-stranded DNA (dsDNA) as the substrate for sample preparation and sequencing. This requirement is seen across a range of platforms, including standalone technologies like PacBio [8], and adapter-mediated methods such as Oxford Nanopore’s nanopore sequencing platforms (MinION, GridION, and PromethION) [9] and Illumina sequencing [10]. Because most sequencing platforms use dsDNA as substrate for sequencing, the protocols detailed in this study for the systematic conversion of ssDNA to dsDNA are extendable to these other sequencing platforms.

**Fig. 1.**
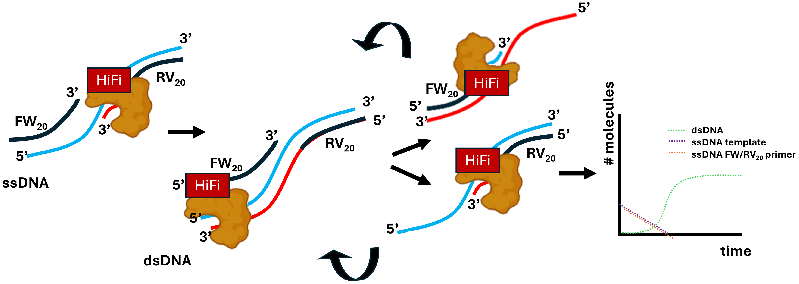
AmpliSeq: Using standard forward (FW_20_) and reverse (RV_20_) primers for exponential amplification of a single-stranded DNA template using PCR. The `HiFi` stands for High-Fidelity, high-temperature, DNA polymerase.

**Fig. 2.**
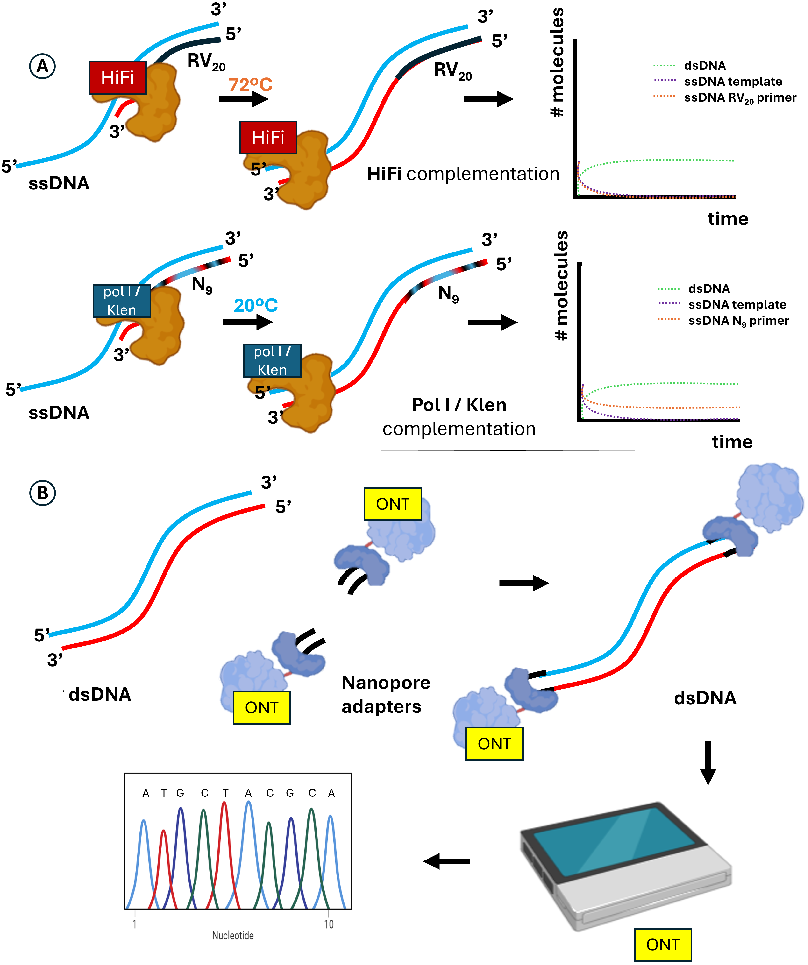
RevSeq: **A)** Using standard reverse primers (`RV`_20_), versus a random primer (nonamers, `N`_9_) approach, for double-stranding complementation of ssDNA molecules enables **B)** ONT-adaptor compatibility and makes the double-stranded substrate suitable for ONT nanopore sequencing readout. `HiFi` stands for High-Fidelity, high-temperature, DNA polymerase (Q5 from NEB, UK) while `PolI/Klen` stands for low-temperature Klenow fragment of the DNA polymerase I.

While different sequencing platforms such as PacBio, Illumina and ONT are available and can be used to replicate the techniques and procedures showcased in this work, which rely on the double-stranding of single-stranded oligonuclotides, the cost-effectiveness of these procedures is what has directed this work towards focusing on ONT technologies (due to its low cost, comparatively) over other sequencing technologies. In Table 1 a comparative analysis of the costs of using ONT, Illumina, PacBio and HPLC/MS is compiled. A descriptive dissection of costs of the aforementioned sequencing platforms is presented in the subsections below to preliminarily justify the use of ONT over other sequencing platforms. Instead, the methods employed for double-stranding, sample preparation and sequencing that have been used for the experiments showcased in this work, using both ONT and Illumina sequencing platforms, are described in the Methods section.

**Table 1.** Cost comparison of different methods for oligo sequencing.

| Method | Minimum Cost<br>(per unit sample) | Double-stranding | Sample<br>prep | Library prep<br>Kit | Barcoding Kit |
| --- | --- | --- | --- | --- | --- |
| ONT | US\$9 | Yes | Yes | Native Barcoding V24 | Included in Library prep |
| PacBio | ≥US\$24.55 | Yes | Yes | Multiple reagents needed | SMRTbell® adapter index plate 96A |
| Illumina | US\$36.99 | Yes | Yes | NEBNext UltraExpress® DNA Library Prep Kit | NEBNext® Multiplex Oligos for Illumina® (96 Unique Dual Index Primer Pairs) |
| HPLC/MS | ≥US\$14.33* (without start-up costs) | No | No | n/a | n/a |

Table 1 catalogues the total sequencing cost per single-stranded oligo using sequencing methods such as ONT, PacBio, Illumina and HPLC/MS. The full-breakdown of the costs is made available in the supplementary section Sequencing costs per platform SS2.

Single-stranded DNA (ssDNA) oligonucleotides are an essential part of the workflows in a molecular biology laboratory, and therefore are valuable for their use in biotechnology, such as in PCR[11], DNA libraries[12], DNA origami[13], and even for DNA data storage[3]. The possibility of sequencing oligonucleotides in a rapid, accessible and standardised manner while still being cost-effective and straight-forward will be a significant achievement in the improving quality control standards of oligonucleotides.

In this light, different strategies are proposed that use both already established and customised techniques for converting ssDNA oligonucleotides into dsDNA, to repurpose currently available and accessible DNA sequencing technology such as Oxford Nanopore Technologies (ONT). ONT technology is used here as a reference sequencing framework, where dsDNA, compatible with ONT adapters, is required for sequencing.

The proposed strategies, which range from PCR amplification, reverse complementation and *random priming* complementation of ssDNA for double-stranding to dsDNA (also referred to as second-strand synthesis) are demonstrated as a means to make an original sample of short ssDNA oligonucleotides into a dsDNA ready for ONT sequencing compatibility. These methods are then compared to the direct sequencing of ssDNA, a negative control incompatible with the binding of ONT adapters and thus unable to yield reads for sequence matching (Figure 3).

**Fig. 3.**
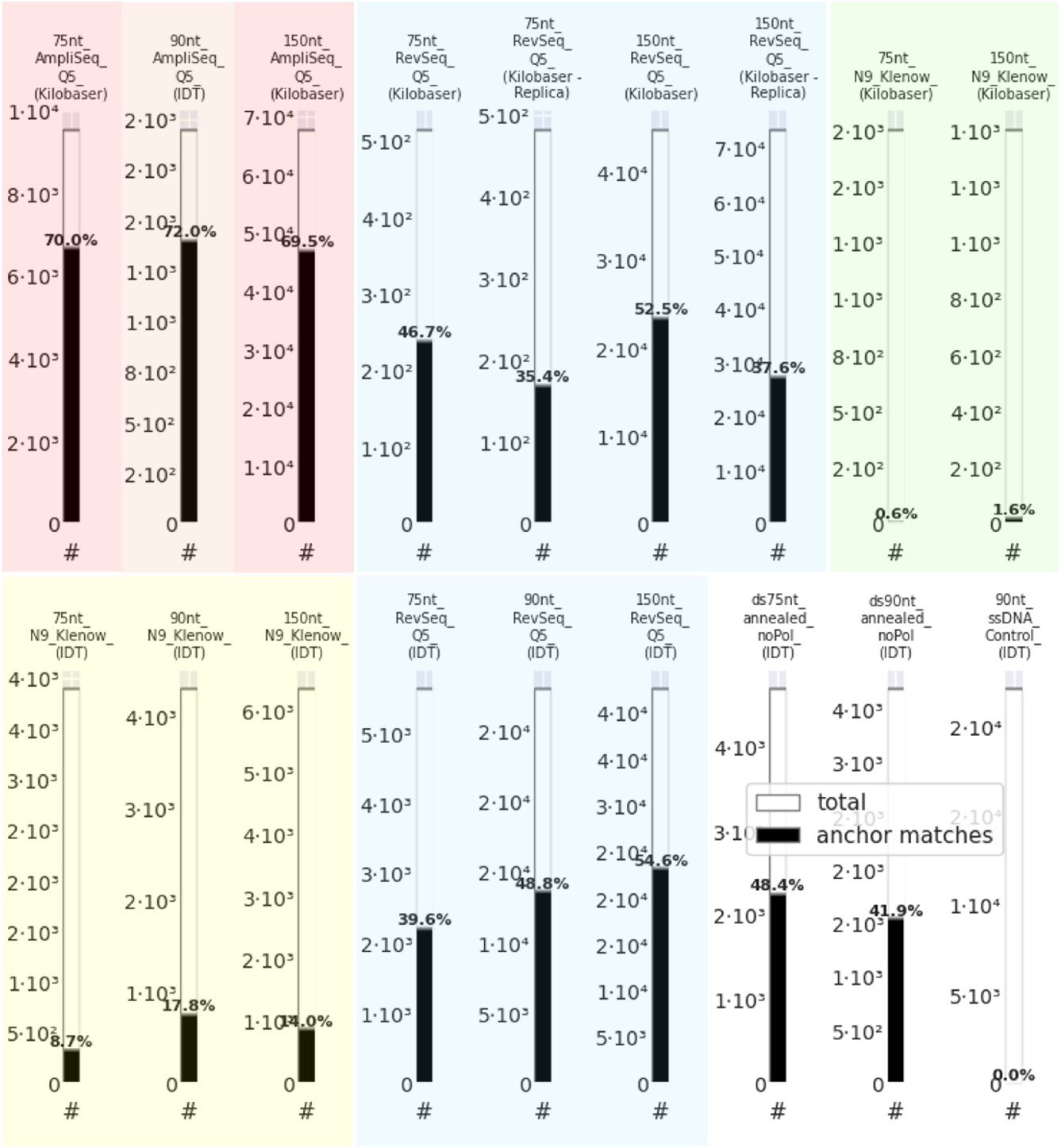
Number (#) of sequence anchor matches that can be aligned to target sequence with ONT sequencing. **Total:** Total number of reads extracted from the sequencing run. **Anchor matches:** Sequences that could be aligned to a reference sequence through anchoring reads to the first 5’ end 20 nt of the plus `(+)` and minus `(−)` DNA sequences filtered using Script S1, regardless of % identity beyond the 20 nt anchors. All experiments were normalised to 200 fmol per load of dsDNA (`AmpliSeq` and `RevSeq`, either with Q5 or N9-Klen) or ssDNA (Control). Substrate ssDNA for all samples was synthesised using an *in-house* Kilobaser or IDT before double-stranding using enzymes or annealing. For a comparison of the results here obtained with those with 50,000 reads using Illumina sequencing see Figure S1.

**Fig. 4.**
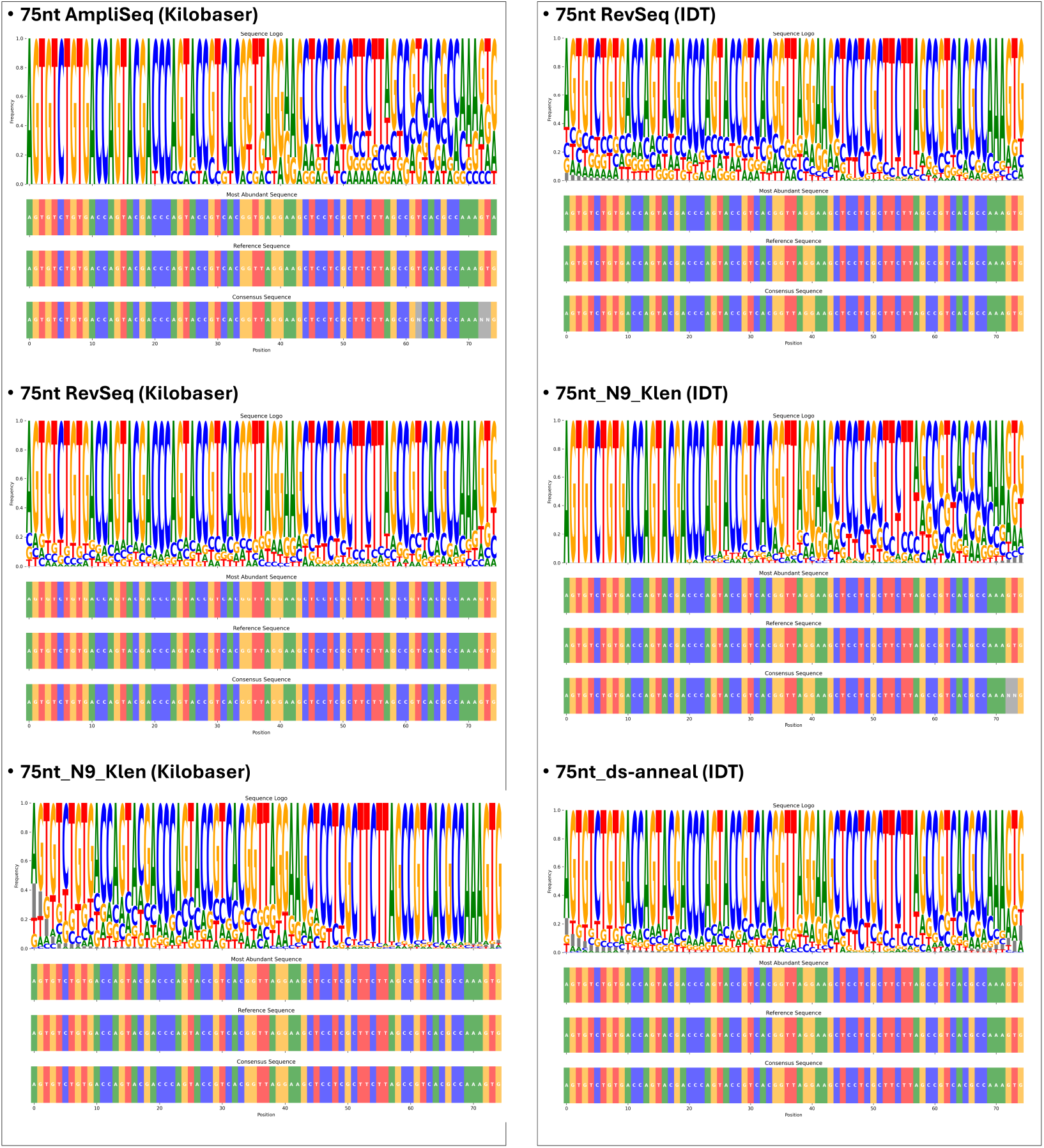
Sequence logo plots for the assessment of the oligo quality of additional 75-mer samples, per sample preparation method. The abundance of a nucleobase at a particular position is proportional to the size of the nucleobase logo in the plot, and the variability is thus represented as a conjunction of sequence logos that can be used to determine a **consensus sequence:** The overall sequence considering all variant reads; and the **most abundant sequence:** which corresponds to the most abundant sequence (and therefore the most likely) after alignment of all reads, when matched against a **reference sequence**.

The number of anchor matching reads with capacity to align (*i*.*e*., sequencing output sequences that can be aligned to the target, or query, intended sequence, see Figure 3) increases significantly when using strategies which make the DNA double-stranded using sequence-specific primer sequences. Therefore, the use of known primer sequences for reverse complementation or PCR-amplification of ssDNA, here termed as `AmpliSeq` and `RevSeq`, (which can be readily extended to RNA sample preparations, despite not shown in this work) into its double-stranded homologous, previous to sequencing processing, yields the best sequencing outputs, *i*.*e*., with the most number of `anchor matches` (Figure 3) that have capacity to positively contribute to meaningful sequence elucidation to the target oligo, or oligo library, sequence able to produce a consensus sequence that is more or less at a pair with its reference (Figure 5). In this work, `RevSeq` is proposed as superior to `AmpliSeq`, despite a decrease in the number of `anchor matches` is observed, because PCR bias may be a concern for some studies and applications where sequence diversification and variability is at stake.

**Fig. 5.**
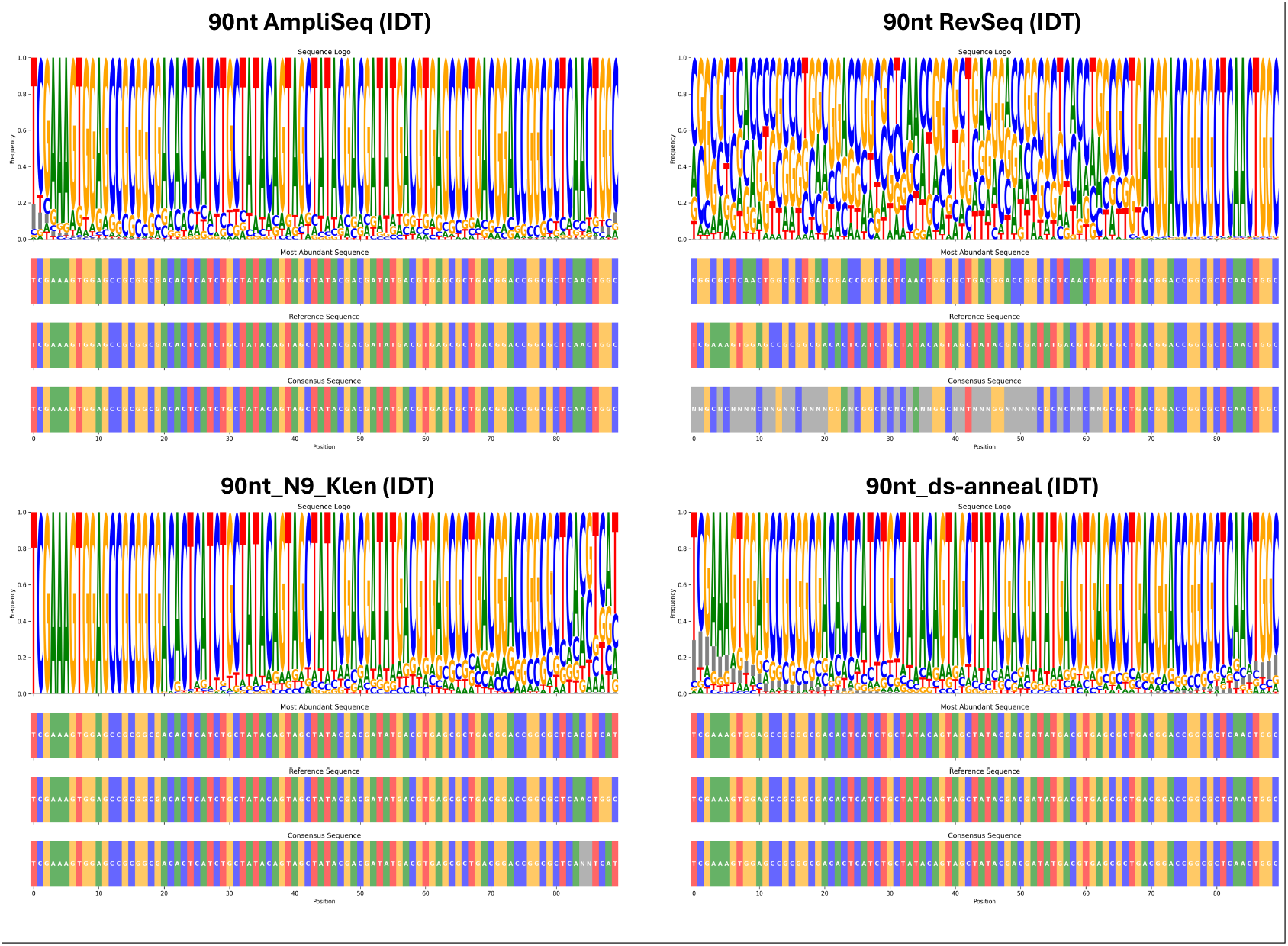
Sequence logo plots for the assessment of oligo quality (90-mer) per sample preparation method. The abundance of a nucleobase at a particular position is proportional to the size of the nucleobase logo in the plot, and the variability is thus represented as a conjunction of sequence logos that can be used to determine a **consensus sequence:** The overall sequence considering all variant reads; and the **most abundant sequence:** which corresponds to the most abundant sequence (and therefore the most likely) after alignment of all reads, when matched against a **reference sequence**.

Overall, the strategies proposed in this work seek to reduce sample processing compared to HPLC/MS while also streamlining the workflow for reverse complementation. By using N9 primers (*N9-Klen* in Figures 3 and 5), our approach eliminates the need for known primers in reverse complementation, which is typically required when using high-temperature polymerases. This aids in the design of simplified pipelines for automation of standardised protocols for studying different samples of ssDNA, which will help, *e*.*g*., with the analysis of stock libraries for the quality control of ssDNA in a rapid and cost-effective manner. This work presents various strategies for DNA sequencing, with the potential to be extended to RNA oligonucleotides using adapter-based technologies such as ONT.

Additionally, these protocols could be adapted for PacBio and Illumina, both of which require a double-stranded substrate for sequencing. By leveraging these approaches, the study eliminates the need for costly quality control methods like HPLC (High-Pressure Liquid Chromatography) or MS (Mass Spectrometry). A key advantage of *OligoSeq* is its ability to provide a rapid, accessible, and comprehensive quality assessment of single-stranded oligonucleotide samples using reverse complementation (Figure 5).

This approach enables evaluation of both read quantity and quality, with read frequencies serving as a reliable guide—a claim validated through two independent methods, `RevSeq` and `N9-Klen` (a modality of `RevSeq` because it relies on reverse-complementation or second-strand synthesis), as demonstrated in Figures 3 and 5. The results obtained highlight that the different methods showcased have different advantages and inconveniences, but that the best approaches necessarily entail the pre-processing of the single-stranded oligonucleotide into its double-stranded equivalent, by either amplification (`AmpliSeq`), reverse complementation (`RevSeq`) (Materials and methods M7), or random primer second strand synthesis using nonamers (`N9`) (Materials and methods M6), prior to sample preparation for sequencing (see Materials and methods M3). The techniques and strategies presented in this work are intended to serve as a valuable reference for both field biologists developing cost-effective sequencing platforms to sequence RNA (*e*.*g*., for transcriptomics analysis) or ssDNA libraries, *e*.*g*., prior to standard cloning, mutant library generation or for highthroughput genome engineering.

### Conversion of single-stranded DNA into double-s-tranded DNA

There are several means to convert single-stranded oligonucleotides (such as DNA and RNA) into its double-stranded equivalent. The omnipresent Polymerase Chain Reaction (PCR)[14][15] is the most readily accessible technique for double-stranding DNA: PCR is a standard laboratory technique that can be used to “amplify” (`AmpliSeq`) or modified to “reverse complement” (RevSeq) short fragments of DNA, including both dsDNA and ssDNA, or RNA, oligonucleotides.

### AmpliSeq: simple PCR-amplification for double-stranding oligonucleotides

The amplification of single-stranded oligonucleotides is possible through the use of short flanking DNA sequences, called oligonucleotides of approximately 20 nucleobases long, that can be used as a “primer” sequences for direct amplification of DNA by means of a DNA polymerase.

Figure 1 demonstrates the basic principle of how two primers (`FW`, *forward*; and `RV`, *reverse*) can be used to direct a DNA polymerase to exponentially amplify a single-strand of DNA after at least a single initial step of reverse complementation (*i*.*e*., Figure 1 shows how the `RV`_20_ primer is first double-stranding ssDNA into dsDNA followed by the subsequent cycles of a standard PCR undergo). Thus, a PCR amplifies the single-stranded DNA strands that have been primed.

While the PCR-amplification procedure shown in Figure 1 above is the most readily accessible and effective (as reflected by the number of anchor matches relative to the other analysed samples in Figure 3) means by which to convert ssDNA into dsDNA, it has the disadvantage that PCR bias is not the most suitable approach to representatively sequence the broad sequence space of *e*.*g*., an DNA oligo library with diversified sequences or point mutations. Likewise, if the goal of *OligoSeq* is the quality control of an oligo sample supposedly having a singular DNA sequence, PCR (`AmpliSeq`) would not be able to representatively capture synthesis errors, degradation, and other quality issues.

In this light, PCR bias refers to the tendency of a PCR-amplified DNA library to lose its sequence diversity during successive amplification cycles, leading to the overrepresentation of certain sequences at the expense of others [16]. This occurs because, in repeated PCR cycles, DNA strands that are amplified earlier tend to become progressively enriched, resulting in their disproportionate dominance within the library. A key strategy to mitigate bias when regenerating DNA libraries is the use of emulsion PCR (ePCR) [11]. This technique, widely utilised in biological research, enables the amplification of a DNA library while preserving sequence diversity. By encapsulating individual DNA molecules within isolated picoliter-scale water-in-oil droplets, ePCR ensures that each sequence undergoes independent amplification, preventing the overrepresentation of subset sequences and maintaining the homogeneity of all possible DNA amplicons in an oligonucleotide library.

### RevSeq: Reverse-complementation techniques for double-stranding oligonucleotides

Reverse-complementation techniques are based on the idea that single-stranded DNA (ssDNA) can be reverse-complemented with a complementary second-strand to form a double-stranded DNA (dsDNA) equivalent that can be suitable for DNA sequencing, such as ONT, but without exponential amplification, and amplification bias, characteristic of PCR; due to only one reverse (`RV`) primer present during the polymerization. The absence of a forward (`FW`) primer in the reaction mixture prevents exponential amplification of ssDNA strands. As a result, only a quantity of dsDNA molecules equivalent to the initial ssDNA substrate remains after reverse complementation, as illustrated in Figure 2. A complementary strand of DNA can be either synthesised using commercially available DNA synthesisers, or by complementing the ssDNA by means of a DNA polymerase (which in this work can be either Q5 (Q5® High-Fidelity 2X Master Mix, NEB, UK) or the Klenow fragment (DNA Polymerase I, Large Klenow Fragment, NEB, UK); and a primer complementary to the reverse flanking end of the complemented DNA sequence (depicted as RV_20_, Figure 2), or a random nonamer (N9) randomly complementary to the reverse-complemented ssDNA (depicted as `N`_9_).

The most evident advantage of using a DNA polymerase to synthesise a reverse complementary strand of DNA, as opposed to synthesising and aligning the same by chemical means, is its cost-effectiveness. A 75-nucleobases long oligonucleotide can cost £42.75 using an external provider (IDT, US) with a 100 nmole DNA Oligo service with a guaranteed yield of 28 nmol or 644.8 *µ*grams (price retrieved from IDT website), or up to £91.68 (excl. taxes) if using an *in-house* oligo synthesiser such as Kilobaser (Kilobaser GmbH, Austria) that needs at least a cartridge (£78.58) having capacity for synthesising 75 Adenines (A), 75 Cytosines (C), 75 Guanines (G) and 75 Tymines (T) and a microfludic chips (£13.10), which usually gives room for synthesis of two oligoes of 75-nt in length, with an expected total yield of approx. 300pmol (recommended oligo length is between 8nt to 50nt according to manufacturer). The cost doubles if a reverse-complementary strand is needed. Instead, with the protocols demonstrated in this work, using a DNA polymerase, such as Q5, Phusion or DNA polymerase I (NEB, UK) the cost can go down to £0.95 per reaction (according to current NEB prices for the reagents involved) which has to be added to the preceding single-stranded DNA synthesis costs.

### Second strand synthesis using high-temperature high-fidelity polymerases

The use of high-fidelity (`HiFi`) polymerases, has the advantage that, by using a known primer sequence of about the standard length of 20-nucleobases (nt), it is possible to reverse-complement oligonucleotides (as depicted in Figure 2) with a representative sample of the errors contained among the synthesised DNA strands (*i*.*e*. avoids PCR bias, as discussed previously).

### Second-strand synthesis using random nonamers

The use of short, random, primers of only 9 nucleobases-long (nonamers) presents itself as a opportunity to mitigate the need to have available a particular primer sequence for the sequence-specific reverse-complementation. Thus, the use of random primers (here nonamers) presents itself as a potential alternative to non-specifically reverse-complement “everything within a DNA library”.

However, the use of random nonamers imposes temperature constraints during reverse complementation, as the priming oligo’s length directly influences its melting and annealing temperatures. Standard PCR reagents, which rely on high-temperature polymerases, are not well suited for this process. As a result, effective enzymatic reverse-complementation requires a polymerase that can function efficiently at lower temperatures. Consequently, standard *HiFi* polymerases commonly used in laboratory PCR are not suitable for applications involving small, random primers. Fortunately, the native DNA polymerase I of *Escherichia coli* is an enzyme that naturally operates at low temperatures of 37 ºC, at its peak performance, and that are suitable for performing at room temperatures (*e*.*g*., 20 ºC on a thermocycler). The protocol used for reverse-complementation is described in Materials and methods M6 and M7.

## Results

### Anchor matches

Figure 3 presents a comparative analysis of sequencing read counts of the different samples processed using `AmpliSeq` (PCR), or `RevSeq` with either a reverse complementing primer (using a Q5 polymerase) or with the random nonamer (N9) approach (using the Klenow Fragment of the DNA polymerase I). For each sample, the total number of sequencing reads (white bars) and the number of reads that passed the anchor filter (black bars), give a measure of the proportion of reads within the total sequencing sample that can be used to sequence an oligo sample, as a way to assess the quality of the procedure to generate a sequencing-able dsDNA (out of the original ssDNA) material of enough quality and representativeness.

As shown in Figure 3, there is substantial variation, across all samples, in both the total number of reads and the proportion of reads that successfully match the anchor criteria (of 20 nt at the 5’ ends of the plus `(+)` and minus `(−)` DNA sequences, respective to a plus `(+)` reference sequence, see Script S1). In several samples, such as the `75 nt AmpliSeq` and `150 nt AmpliSeq Q5 (Kilobaser)` and the `150 nt RevSeq Q5 (Kilobaser - Replica)`, a high proportion of reads (ranging from approximately 52% to 70%) pass the anchor filter (see Materials and methods M5), indicating efficient and specific amplification (or reverse complementation in case of `150 nt RevSeq Q5 (Kilobaser - Replica)`) or sequencing of the target region. This suggests that the use of both PCR and reverse complementation with defined primers using a Q5 polymerase was effective in generating double-stranded reads that align well with the reference anchors (without regard to the rest of the sequence, which is analysed in Figure 5).

Conversely, other samples, such as the `75 nt` and `90 nt N9 Klenow (Kilobaser)`, show a much lower proportion of anchor-matched reads (as low as 0.6% to 1.6%), despite having a moderate total read count (*i*.*e*. all samples were normalised to 200 fmol as evidenced with the consistence of the total reads accross samples in Figure 3). This low efficiency could reflect issues such as non-specific reverse complementation or amplification using random non-amers (`N9`), or a high background of off-target sequences due to the origin of the single-stranded DNA. However, because when using quality samples synthesised with IDT (see `75 nt` and `90 nt 150 nt N9 Klenow (IDT)` gave an over two-fold (logarithmic) improvement in the number of anchor matches (Figure 3) with values of 8.7%, 17.8% and 14%, respectively, it is fair to suggest that quality of the original sample of DNA, rather than unspecific or offtarget amplification, is an observable feature of the results obtained with an *in-house* synthesised single-stranded DNA (Kilobaser) versus an industry-standard single-stranded DNA (IDT).

As expected, with the `90 nt ssDNA Control (IDT)` (used as a negative control), no reads passed the anchor filter because no double-stranding of single-stranded DNA was realised.

Intermediate results were observed in samples like the `75 nt` and `90 nt RevSeq Q5 (IDT)`, where approximately 39–49% of reads are anchor-matched, which may suggest partial success in targeting the desired region, but the authors attribute it to the capacity of `RevSeq` to capture the variability of the oligonucleotide sample to a much greater degree than the `AmpliSeq` samples, because `RevSeq` avoids PCR bias and therefore is able to show greater variability within a sample sequence (as depicted in Figures 4—6).

The `dsDNA annealed noPol (IDT)` samples (for `75 nt` and `90 nt`, respectively) were used here as a positive control in which no enzymatic processing of singlestranded DNA was realised and instead the reverse complementing DNA strand was provided and annealed to the leading strand as a means to obtain the dsDNA intermediate for ONT sequencing sample preparation (Materials and methods M3), using industry-grade DNA from IDT. This positive control shows that, when using the dsDNA-annealed IDT samples, moderate anchor-matching rates of up to 41–48% can be expected, indicating that a significant fraction of the reads correspond to the intended sequence. These controls also establish that when using ONT, it is expectable to see a consistency of over 50% of total sequenced reads not being meaningful to the reference sequence. This low percentage of meaningful reads can attributed to the debris found from DNA synthesis (for both IDT and Kilobaser samples) that come naturally from errors or aborted synthesis due to capping, in phosphoramidite synthesis, that ultimately lead to a heterogeneous sample of more or less quality depending of the methods of purification. However, methods such as purification using agarose gel electrophoresis, despite could be used to clean-up incomplete sequences below the desired length, would add up to the cost of synthesis.

### Sequence logos

Despite sequence variability and the presence of incomplete sequences that could be avoided with gel purification, the full sequence can still be retrieved correctly based on both the “most abundant sequence” and “consensus sequence” as exemplified in Figures 4—6. Surprisingly, synthesising both strands at industry quality grade (i.e. using IDT as an external provider) did not yield better performance (measured as high `anchor matches`, Figure 3, and low frequency variability, Figure 5, in the `ds-anneal` bottomright plot) than the observed with `AmpliSeq`, further reinforcing the idea the synthesis quality can be evidenced using this protocol, and that the remarkably better results of the `AmpliSeq` samples, are likely due to PCR bias and the enrichment of meaningful reads due to amplification of a more homogeneous anchor matching sequence pool, which does not occur with the `RevSeq` samples, nor with the directly annealed dsDNA samples, because these better reflect the variability of sequences implicit of samples with high, but still limited to a certain extent, synthesis quality.

Remarkably, the rationale that AmpliSeq gives significantly larger number of anchor matches, respective to RevSeq, due to RevSeq double-stranding only the diversity of sequences available for second-strand synthesis, while AmpliSeq can amplify the pool to as many sequences as primers are available, are not corresponded when using the 50, 000 reads extracted from “sequencing-by-synthesis” Illumina’s technology (Figure S1), which is used as comparison to ONT sequencing to determine to which extend the results obtained can be influenced by sequencing technology. Although it is clear that these differences can be attributed primarily to the differences in sequencing technology (e.g. sequencing by synthesis in Illumina, using clusters and reading averages, versus the single-molecule read capability of ONT), the authors refrain to speculate on what the particular causes can be involved in these differences, since the sample preparation steps between ONT and Illumina are very similar and all samples were normalised to 200 fmol for sample preparation.

Overall, these results demonstrate the effectiveness of the methods proposed for the rapid sequencing and quality control of single-stranded oligonucleotides using nanopore sequencing and highlight the importance of sequencing depth and the specificity of read alignment to the reference anchors (*e*.*g*. high total read counts are only valuable if a substantial proportion of those reads are relevant to the target region). The anchor-matching percentage (Figure 3) serves thus as a key quality metric, reflecting the specificity (and bias) of the experimental protocol for each sample (*i*.*e*. accounting for `AmpliSeq` or `RevSeq`, specificity of primers and polymerases).

Figures 4—6 present sequence logo plots for each sample (75-, 90- and 150-mers) and double-stranding method (AmpliSeq and RevSeq, with or without random N9 primers), illustrating the nucleotide frequency at every position along the synthesised DNA oligonucleotides. These plots provide a detailed view of sequence conservation and variability across the different synthesis and reverse complementation strategies. Each logo comes with a colour-coded representation of the most abundant sequence, the reference sequence, and the consensus sequence, to facilitate a direct visual comparison. Across most samples, the sequence logos reveal high conservation at the anchor regions, as expected from the anchor-filtering approach described in Figure 3. However, it is important to highlight that the nature of alignment, because it starts with the sequences that were anchored also conditions a higher conservation of sequences at the anchor sites with the most abundant sequences. This can be reflected by taller and less diverse defined letters at a particular position at the sequence ends (Figures 4—6), indicating that the majority of reads align more precisely to the reference at these positions. In contrast, the more central regions of the sequences may display varying degrees of nucleotide diversity, with some samples showing a more pronounced sequence variability as evidence of mutations or synthesis errors. Although this anchoring bias is not visibly evident in most of the samples characterised in Figures 4—6, the anchoring effect should still be considered to understand displayed errors. This is more patent by the fact that DNA samples with longer lengths e.g., 150-mers (Figure 6), naturally display a greater anchoring effect than its shorter counterparts (i.e. 75- and 90-mers) due to the greater proportion between anchors (20-nt at both ends) and the rest of the sequence. For example, samples synthesised using the N9 Klenow or ds-anneal approaches (notably those from IDT) show increased sequence heterogeneity, as indicated by shorter, more mixed stacks of letters in the logo plots. This suggests a higher frequency of incorporation errors or incomplete synthesis, particularly in the regions in between the 20-nt flanking anchors of the analysed oligonucleotides.

**Fig. 6.**
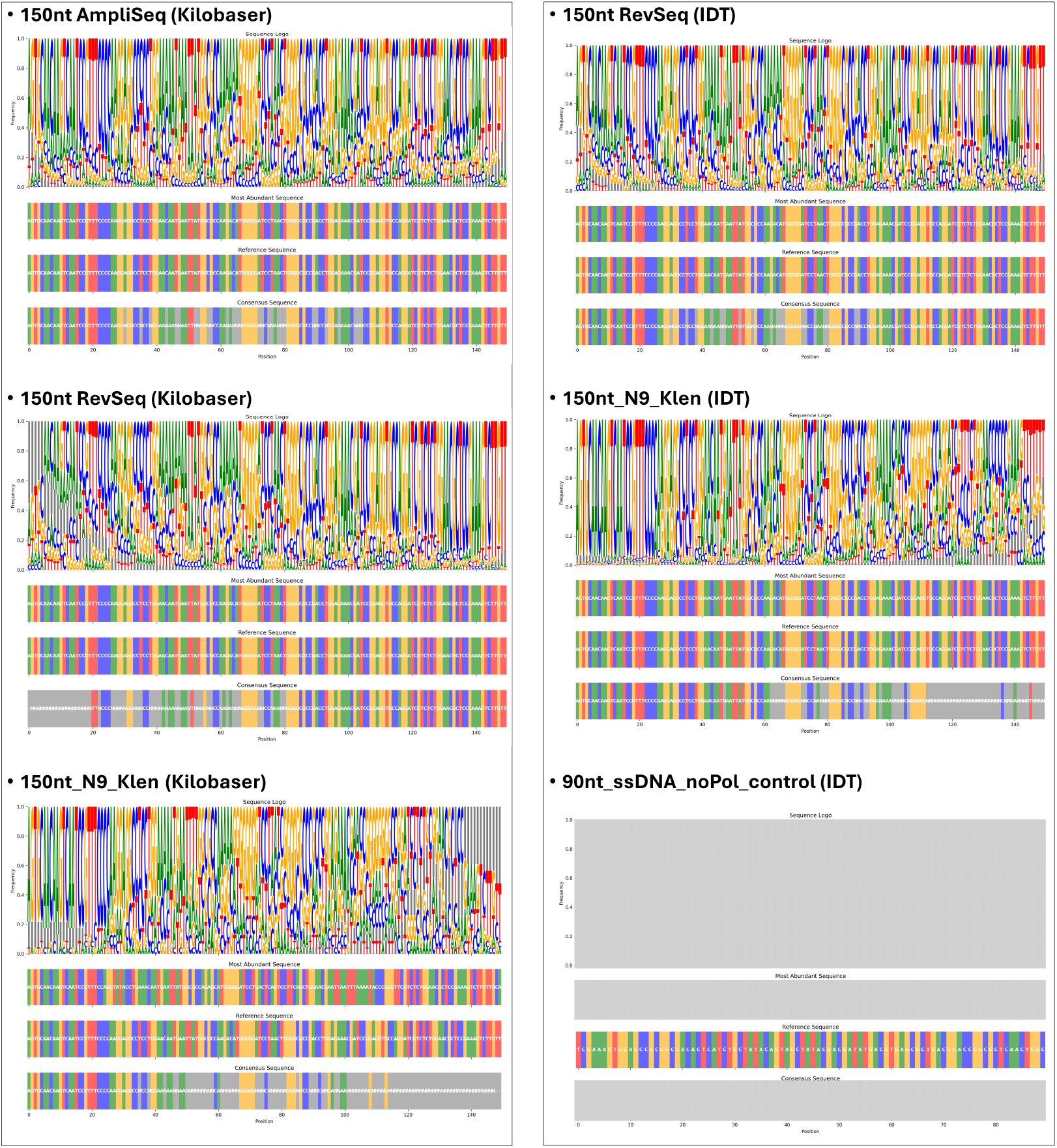
Sequence logo plots for the assessment of the oligo quality of additional 150-mer samples, including a single-stranded 90-mer blank control, per sample preparation method. The abundance of a nucleobase at a particular position is proportional to the size of the nucleobase logo in the plot, and the variability is thus represented as a conjunction of sequence logos that can be used to determine a **consensus sequence:** The overall sequence considering all variant reads; and the **most abundant sequence:** which corresponds to the most abundant sequence (and therefore the most likely) after alignment of all reads, when matched against a **reference sequence**.

In regards to the frequency of deletions displayed in Figure 5, it is important to justify the fact that deletions at the ends of the sequences do not necessarily imply an accumulated frequency of deletions at the end positions, but that these deletions can accumulate anywhere in the sequence and then evidenced as deletions at the end-sequence due to the shortening of the aligned sequences to the reference sequence (which in Figure 5 are always aligned without penalisation for deletions, substitutions or insertions due to the purpose of the plots simply being the evidence of sequence variability, rather than alignment of sequences to a reference). Therefore, the observed deletions at the end-sequences contribute to the frequencies observed at each position throughout the entire sequence, as a result of shortened sequences in case of point deletions (at any point of the sequence) that produce the frame shifts, e.g., `150nt N9 Klen` (Figure 6).

Therefore, these “shortening” (also mentioned as incomplete) of sequences due to synthesis problems (*i*.*e*. insertions, deletions or substitutions) can be attributed to both, deletions at any position within the sequence, and termination of synthesis due to extended lengths (which is more patent when using Kilobaser 150-mers, respective to their IDT counterparts). This observed sequence variability due to termination of synthesis is also consistent with the chemical synthesis process, which proceeds from the 3’ to 5’ end of the plus `(+)` strand. Errors and mutations tend to accumulate toward the 5’ end, as each successive nucleotide addition carries a risk of incomplete coupling or side reactions. Additionally, the subsequent reverse complementation, performed using various enzymatic strategies, further influences the fidelity of the final double-stranded product. In this regard, samples prepared with the high-fidelity polymerase (Q5) and biasing protocols (*i*.*e*., `AmpliSeq`) generally display lower sequence variability and fewer mutations, as evidenced by the dominance of the reference base at each position in the logo plots. In contrast, samples subjected to less stringent, more diversely representative, or more error-prone reverse complementation `e.g.` using either the `RevSeq` with normal reverse-complementing primers, or a random nonamer (N9) for the Klenow fragment of the DNA polymerase I, show greater sequence diversity, with some positions showing no clear consensus base. In the same line, sequences synthesised using industry grade DNA, particularly at DNA lengths longer than 75 nt also display improved metrics (i.e. a decreased diversity of sequences and a more consensus sequence).

Overall, and despite minor differences that can be attributed to PCR bias, substrate DNA length and quality of the original DNA sample, it is possible to observe that all studied cases are able to retrieve fully matching sequences (with degree of variability implicit to sample quality), and where consensus is lesser (particularly observed at DNA sequences of 150 nt) it is still possible to retrieve the original sequence using the “Most Abundant Sequence” based on the most frequent aligned sequences retrieved from the sequence logo of Figure 6. These findings are in agreement with the anchor-matching results shown in Figure 3, where samples with high anchor-matching rates also tend to show greater sequence fidelity throughout the oligonucleotide. Conversely, samples with low anchor-matching rates often display increased sequence heterogeneity and a higher incidence of mutations, reflecting both synthesis and processing inefficiencies. Notably, the `90 nt ssDNA noPol control` sample shows no meaningful sequence logo, consistent with the absence of anchor-matched reads in this condition.

Overall, the sequence logo analysis underscores the impact of both chemical synthesis and enzymatic processing on the quality and fidelity of synthetic DNA oligonucleotides. The combination of anchor-filtered read selection and sequence logo visualization provides a useful approach for assessing sequence integrity, identifying sources of variability, and optimizing protocols for high-fidelity DNA synthesis and processing.

The analysis of Figure 3 shows that `AmpliSeq` is the method that yields the most homogeneous sample, with the highest proportion of aligning reads that fulfil the proposed cut-off anchor sequences of 20 nt at both 5’ and 3’ ends of the reference sequence (Figures 4—6). This is consistent with the observation that PCR can enrich the sample of ssDNA into an exponentially amplified dsDNA sample, but also of PCR bias. Instead, the results obtained for the `RevSeq` and N_9_ experiments *i*.*e*., either with `HiFi` Q5 polymerase, or the Klenow fragment of DNA polymerse I, show much lower number of target anchor reads, which belong to the fact that reverse-complementation does not amplify the sample, but simply produces the complementary strand of the leading strand of ssDNA, thus producing samples with similar number of molecules at the input than at the pre-processing output (*i*.*e*., input ssDNA vs dsDNA output), which according to the positive controls, should be no higher tan 50% (Figure 3). However, within reverse-complemented samples, a consistent, orders of magnitude, decrease in the relation of the number of total and target anchor match reads, between all samples is obtained. An increase in the variability of the anchor matching target sequences from the oligo sample is also observed, respective to results obtained with the PCR-based `AmpliSeq` experiments, which is consistent with the hypothesis of PCR bias mentioned in the introduction, and the fact that the nature of reverse-complementation does circumvent it.

The two different methods for reverse-complementation used, RV_20_ and N_9_, show that similar variability profiles (as demonstrated in Figure 5) are obtained, despite a remarkably lower proportion between anchor matches and total reads, with the advantage that using N_9_ random nonamers facilitate not having to continuously pre-design and re-design primer sequence and give room to a more simplified protocol that is more amenable to automation and high-throughput processing, due to the fact that primer design is not needed for reverse-complementation with random primers.

In this sense, `RevSeq` is presented here as a useful method that enables the sequencing of input samples of ssDNA that can be heterogeneous and therefore have a greater variability, needed in experimental set ups where large and diverse DNA and RNA libraries are used (*e*.*g*. a library of mutants), through a protocol that is accessible, rapid, and standardisable, using either `HiFi` polymerases with standard, 20-nucleobase long, primers, or the use of N_9_ primers (Figure 2) as an interesting alternative for quality control of oligonucleotide samples.

The `RevSeq` approach, with its RV_20_ or N_9_ variants, demonstrated has the advantage that the PCR amplification bias can be circumvented, despite at the expense of a lower number of `anchor match` reads (Figure 3) that can be aligned to a reference sequence based on 20 nt anchors (at both sides of the reference sequence, Figures 4—6), and is therefore presented as a suitable approach for determining variability within a sample library of oligonucleotide sequences which, as aforementioned, could comprise diversified point mutations, due to randomization, variant selection, or oligo synthesis quality issues.

Using the results demonstrated in Figures 4—6 it is easy to see that when using a challenging sample e.g., a Kilobaser synthesis of a 150-mer (`150nt_AmpliSeq`, `150nt_RevSeq` and `150nt_N9_Klen`, Figure 6), the oligonucleotides analysed display a greater diversity and a lack of consensus, which although does not prevent to retrieve the original sequence using “the most abundant reads”, exemplify the limitations of synthesis of long oligonucleotides (including industry-grade standard 150-mers from IDT) compared to more feasible sequence lengths (75-mers and 90-mers) also tested in this work. In this work, a cut-off sequence of 75 nt-long was used, taking into account that, for long, industry standard single-stranded oligonucleotides have been capped to 80–90 nucleobases.

In this line, for long 150-mers, the origin of the oligo sample decreases significantly in the Kilobaser synthesised oligos, respective to the IDT synthesised ones, despite the variability (upper frequency plots in Figure 6) shows that always the most likely sequence matches that of the query reference used for synthesis in all cases. This should of course be attributed to the differences in sample quality that come from a different procedure for synthesis and purification and the quality control checks in place, which are patent in IDT samples, but not in our procedure for oligo synthesis using Kilobaser.

Remarkably, for 150-mers where full DNA lengths are more difficult to achieve (synthesis-wise), the general increase in frequencies usually show a slight increase in the variability of sequences a the 5-end of the sequenced oligonucleotides, which translates into a consensus sequence position labelled as “`N`”, which is consistent with the fact that sequences are synthesised 3’ end to 5’ end. However, errors at the sides of each frequency plot should also be attributable to the nature of the alignment that always goes from the anchor to the rest of the sequence, prioritising better reads early in the alignment. To mitigate this artifact, both references (plus `(+)` and minus `(−)` strands) are used for the alignment (see Script S1 and Figure 7).

**Fig. 7.**
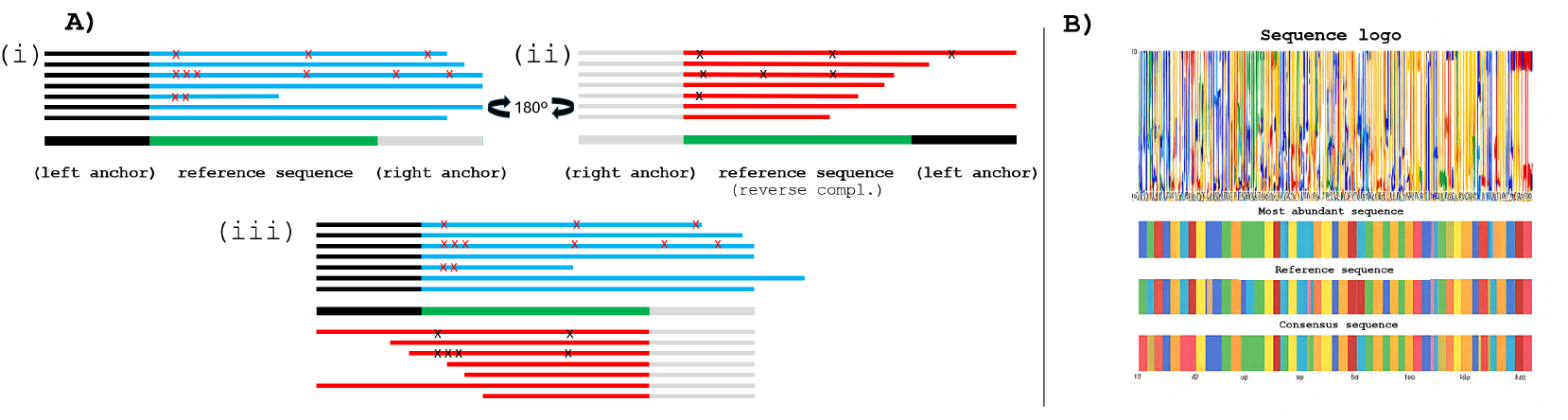
Strand-aware alignment and sequence logo generation using anchor-based read filtering. **A)** Schematic representation of anchor-based alignment: (i) Forward reads are aligned based on matching a predefined left anchor (black), extending into the reference sequence (green) and right anchor (gray); mismatches (mutations) are marked with red crosses (x). (ii) Reverse reads are reverse-complemented and aligned based on matching a right anchor (black, right), proceeding leftwards along the reference. (iii) All reads passing anchor filters—forward and reverse—are aligned and combined into a uniform forward sequence dataset. **B)** Output visualization of the aligned reads: the sequence logo (top) represents nucleotide frequencies at each position; below, the most abundant sequence, reference sequence, and consensus sequence are shown using colour-coded nucleotide highlights for intuitive and direct visual comparison.

In summary, from the analysis of the results demonstrated in Figures 4—6, when using an oligonucleotide sample synthesised by IDT (Integrated DNA Technologies, US) *i*.*e*., the `90nt_RevSeq` (using Q5 and a RV_20_ primer) vs `90nt_N9_Klen` (using the Klenow fragment of the DNA polymerase I and a N_9_ primer) show consistent results in regard to the variability displayed for the different reads of the sample. This highlights that both `RevSeq` strategies (RV_20_ vs. N_9_) are equally valid and demonstrate a similar variability which cannot but be correlated to the real diversity of sequences within the analysed sample. The results also demonstrate that what is described for the 90 nt sequence (synthesised and quality-guaranteed by IDT) does also apply to the *in-house* Kilobaser-synthesised samples of 75 nt and 150 nt, respectively, which display similar profiles for the two methods (RV_20_ vs. N_9_) of `RevSeq` used. This is also indicative that the improvement in the number of reads; but more importantly, the increased (although not remarkably) sequencing quality (as exemplified by a less diverse sample and a more defined consensus sequence of the samples processed using `AmpliSeq` (*i*.*e*. PCR) is likely to be due to PCR bias.

In regards to the 90 nt sequence (IDT), it is observed, that for the cases of reverse complementation, the yield of high identity matching sequences was slightly lower (Figure 3, without a compromise in the final sequences retrieved through “the most abundant sequence” shown with its variability (Figure 5). This can be attributed to the fact that the 90 nt sequence showcased, comes from an IDT tube that over-a-year-old (compared with the other samples that were recently synthesised) and that has been frozen and thawed multiple times throughout this year, while all other samples where immediately processed after synthesis. This highlights the more how the techniques shown in this work are of value to assess the quality of single-stranded oligonucleotide samples, which can be compromised by synthesis technique but also by storage conditions and time.

Ultimately, Figures 4—6 demonstrate how `RevSeq`, and specially with the random nonamers (N9) technique, can be of use for preliminary quality control of single-stranded oligonucleotides that span from 75 nt to 150 nt. Of relevance is the consistency to which the reverse complementation, either using Q5 polymerase or the Klenow fragment with random nonamers, is able to represent the variability of the sample. In this sense, the consistency shown when comparing of Q5 and N9 is of interest because using random primers (*i*.*e*. N9 nonamers) allows to automate the process of quality control without regard to the sequences to analyse, and with the possibility to do this at room temperatures (*e*.*g*. 20 ºC) which is useful for field applications outside the laboratory.

The results demonstrate a first-in-kind sequencing and quality control protocol of single-stranded oligonucleotides using a DNA sequencing technology (in this case using ONT, but the sample processing could be certainly adapted for PacBio and Illumina, despite at a larger cost and processing times), as an alternative to more comprehensive (and expensive) methods such as HPLC/MS.

#### Comparative analysis between ONT and Illumina sequencing

The main results demonstrated for OligoSeq rely on ONT/nanopore sequencing and focus on showcasing different techniques that can be used to adapt single-stranded oligonucleotides to the available sequencing technologies through double-stranding (i.e. using AmpliSeq or RevSeq, either with sequence-specific primers or random nonamers). Therefore, in an attempt to assess also the sequencing accuracy of the methods used, and to provide experimental information about how the sequencing platform of choice (i.e. ONT or Illumina) can affect the double stranding methods here shown, this study also shows Figure S1 and Figure S2 as a comparison between the different double-stranding strategies, but validating the performance of nanopore sequencing (ONT) versus Illumina sequencing experimental results using the same samples amplified using the methods for double-stranding here showcased.

Therefore this work results in the first comparative analysis between the performance of DNA sequencing using ONT versus Illumina for short oligonucleotide sequencing using the methods for double-stranding, sample preparation and sequencing shown in this work. Figure 3 and Figure S1 show the percentage of anchor matches obtained between ONT and Illumina sequencing technology. For the Illumina read samples, a 50, 000 reads was extracted from a sequencing run, in order to normalise the number of reads to a level similar to those reads obtained when sequencing 200 fmol of sample as recommended by the ONT sample preparation protocol.

As seen when comparing these figures, the results obtained show that the proportion of reads with ≥90 % anchor matches is inferior in all cases, and remarkably inferior with the AmpliSeq samples, which fall more than 50 % respective to the results obtained using ONT sequencing. The results from RevSeq show that 29.4 % seems to be an upper limit of anchor matches which can be attributed to the fact that the proportion of “noise reads” using Illumina is always greater than that obtained when using ONT sequencing. The results obtained using N9_Klenow as a double stranding strategy also show a smaller amount of *≥*90% anchor matches when using Illumina over ONT, highlighting once again the superiority of ONT over Illumina for the sequencing of oligonucleotides using these double-stranding approaches, particularly when using the random nonamers method (i.e. N9_Klenow).

Figure 4, Figure 5 and Figure 6 show the sequencing logos for ONT sequenced oligoes of 75-, 90- and 150-nt, respectively, and Figure S2 shows the comparative frequencies corresponding to the sequence logos obtained when using Illumina.

In regards to the sequencing technology used, the comparison between Figure 3 and Figure S1 highlights that using ONT technology, over e.g. Illumina, does not compromise (but rather improve) the proportion of anchor match sequences that can be aligned to resolve the final sequences using the sequence logos of Figure 4, Figure 5 and Figure 6, respectively for ONT, versus Figure S2 for Illumina. These results highlight how this greater abundance of matching reads, also with greater variability respective to the reference sequence, can be attributed to the differences between ONT and Illumina sequencing technology: while ONT produces as many reads as actual molecules are present in the sample, Illumina relies on clustered sequences using sequencing by synthesis technology, which may contribute to a decrease in the number of anchor matches (Figure 3 vs Figure S1) and also a decrease in the varaibility observed in the sequence logos, in which a decrease in the number of matches per every 10^4^ reads depends (Figures 4—6 vs Figure S2).

## Conclusion

As expected, the results obtained highlight `AmpliSeq` (PCR), and `RevSeq` (reverse-complementation) as methods for the sequencing of single-stranded DNA oligonucleotides of 75 nt, 90 nt, and 150 nt in length. For all cases, a correspondence between an increase in length and a decrease in quality is observed, regardless of whether the oligos were synthesised using Kilobaser or IDT, although the quality was visibly better when using IDT. A decrease in the quality of IDT samples when using oligos that have been stored and re-stored for over a year (with repeated freeze-thaw cycles) is also observed, respective to samples recently synthesised. A correlation between increased length and lower qualities (as corresponded by increased sequence diversity and decreased frequencies) is also observed as a measure that can help define quality which can be found when using `RevSeq` (either with Q5 or N9-Klen), as opposed to the more prone-to-bias `AmpliSeq`.

In regards to the comparison of the sequencing method used after double-stranding of single-stranded oligonucleotides, the results show significant differences in the number of anchor matches with ≥90 % identity obtained when using as many reads as possible with ONT (overall 10^4^ reads) versus the 50, 000 extracted from the pool of obtained reads coming from Illumina. These results show a greater number of anchor matching reads for ONT versus Illumina (Figure 3 vs. Figure S1). However, the lower number of anchor matches for Illumina do not prevent to resolve the final sequence using the sequence logos shown in Figure S2 for Illumina, versus the ones shown in Figures 4—6 for ONT and the results obtained with Illumina show a much lower varaibility of sequences, which may come from the fact that ONT is more prone to substitutions and indels than Illumina due to its nanopore-based single-molecule read technology. These results, adding up to the fact that, as shown in Table 1, the costs per run of ONT are far more cost-effective that any other technology here studied, justify the use of ONT technology for the rapid sequencing of single-stranded oligonucleotides.

The results demonstrate that Illumina, while displaying somewhat generally “cleaner” sequence logos (Figure S2) and in the particular case of RevSeq for 90nt oligo, a non-erroneous better sequencing output, does not show a superior sequencing output, respective to the results obtained when using ONT sequencing, as exemplified by the fact that all sequence logos obtained using ONT (Figures 4—6) show a resolved final sequence that matches to the originally sythesised one (either with Kilobaser, or from the external provider IDT), to the exception of the 90-mer sample that has undergone RevSeq for double-stranding (Figure 5).

Overall, the results in Figure 3 and Figure 5 show that both `AmpliSeq` and `RevSeq` are appropriate techniques for the pre-processing of single-stranded oligonucleotides for oligonucleotide sequencing (*OligoSeq*) using ONT that are presented as a rapid and accessible alternative to currently standardised oligonucleotide quality control via HPLC and/or MS; while `RevSeq` is presented as a method that is able to show a greater variability, which increases as the synthesis quality of the oligo sample decreases, demonstrating the potential of `RevSeq` to study sequence diversity within an oligo library for many applications in biotechnology, such as oligo quality control, assessment of the representativeness of a diverse ssDNA pool, or for mRNA sequencing using cDNA libraries, among many others.

## Materials and methods

The materials and protocols used for the preparation of this manuscript are stated as follows.

### M1. DNA sequences

The following DNA sequences were synthesised using IDT (US) or Kilobaser GmbH (Austria) for use in the experiments shown in this work.

**Script 1**. DNA sequences used in this work.

~~~
>75nt
AGTGTCTGTGACCAGTACGACCCAGTACCGTCACGGTTA
GGAAGCTCCTCGCTTCTTAGCCGTCACGCCAAA
GTG
>75nt_FW_primer
AGTGTCTGTGACCAGTAC
>75nt_RV_primer
CACTTTGGCGTGACGG
----------------------------------------
>90nt
TCGAAAGTGGAGCCGCGGCGACACTCATCTGCTATACAGT
AGCTATACGACGATATGACGTGAGCGCTGACGGACCGGCG
CTCAACTGGC
>90nt_FW_primer
TCGAAAGTGGAGCCGCGGCG
>90nt_RV_primer
GCCAGTTGAGCGCCGGTCCG
----------------------------------------
>150nt
AGTGCAACAAGTCAATCCGTTTCCCCAAGGAGGCCTCCTG
GAACAATGAATTATGGCGCCAAGACATGGGGGATCCTAAC
TGGGGCGCCGACCTGGAGAAACGATCCGGAGGTGCCAGGA
TCGTCTCTGGAACGCTCCGAAAGTCTTGTT
>150nt_FW_primer
AGTGCAACAAGTCAATCCGT
>150nt_RV_primer
AACAAGACTTTCGGAGCGTT
~~~

#### M2. Selection of experimental samples for analysis

Sequences of 75 nt, 90 nt and 150 nt in length were chosen to highlight higher error rates positively correlated to the length of the oligos used in this work. The samples used were synthesised by either a DNA Express machine (Kilobaser Gmbh, Austria) or Integrated DNA Technologies (IDT) as a means to have a control samples with an industry standard quality control (*i*.*e*. IDT). `AmpliSeq` (PCR), and `RevSeq` (reverse-complementation) of single-stranded DNA oligonucleotides of 75 nt, 90 nt, and 150 nt in length were realised as a means to compare the performance of the different proposed methods with the length of the oligonucleotides used. When using Kilobaser, a correspondence between an increase in length and a decrease in quality should be expected, and a control using oligonucleotides synthesised by IDT is thus provided, showing also a similar correlation between increased length and lower qualities (as corresponded by increased sequence variability and decreased frequencies) and to assess whether an increase in frequency diversity is found when using `RevSeq`, as opposed to the more prone-to-bias `AmpliSeq`. A control run consisting of an unprocessed, industry-grade, quality controlled, single-stranded 90-mer sequence from an external manufacturer (IDT, US) was used.

#### M3. ONT barcoded sample preparation

All DNA oligonucleotide preparations were resuspended in nuclease-free water (for Kilobaser-synthesised samples) or in IDT-buffer (for IDT control samples) and double-stranded using either `AmpliSeq` or `RevSeq` (with either the RV_20_ or the N_9_ primers), the resulting amplified or reverse-complemented products were cleaned-up with two consecutive steps of 80% EtOH, after mixing the sample with a 180 %(v/v) ratio of AMPure XP beads (Beckman, US). Next, a DNA dA-tailing procedure was effectuated using NEBNext® Ultra™ II End Repair/dA-Tailing Module (NEB, UK) by mixing 11 *µ*L with 200 fmol of oligonucleotide sample with 1.75 *µ*L of NEB-Next® Ultra™ II End Repair/dA-Tailing Buffer and 0.75 *µ*L of NEBNext® Ultra™ II End Repair/dA-Tailing Enzyme Mix, and incubated at 20°C for 5 minutes and 65 °C for an additional 5 minutes before purification again with AMPure XP beads. A 7.5 *µ*L of each end-prepped doubly-stranded sample was mixed with 2.5 *µ*L of a different Native Barcode (NB01-24), 10 *µ*L of Blunt/TA Ligase Master Mix (NEB, UK) before incubation for 20 minutes at room temperature. The each barcoding reactions was stopped with 4 *µ*L of a solution of EDTA (SQK-NBDD114.24 - ONT, UK) before pooling all reactions into a same Eppendorf tube. The resulting pool was mixed with a 40 %(v/v) ratio of AMPure XP beads (Beckman, US) and cleaned-up with two consecutive steps of 80% EtOH before eluting a mixed pool in 35 *µ*L nuclease-free water. A ligation reaction by combining 30 *µ*L of the purified end-repaired and dA-tailed pool of oligonucleotides was mixed with with 10 *µ*L of NEBNext Quick Ligation Reaction Buffer (5X), 5 *µ*L of Native Adapter (SQKNBDD114.24 - ONT, UK) and 5 *µ*L of Quick T4 Ligase (NEB, UK) incubated for 20 minutes at room temperature. After ligation, a successive modified step for AMPure XP cleanup, in which instead of the two steps of 200 *µ*L of 80 % EtOH for paramagnetic-bead clean-up, two-steps of 125 *µ*L of Short Fragment Buffer (SFB) (SQK-NBDD114.24 - ONT, UK) is used, to remove excess adapters. The DNA is then quantified using a suitable method, such as a fluorometer. Finally, the prepared DNA are loaded onto the nanopore flow cell or flongle, according to the manufacturer’s instructions, ensuring proper settings on the MinION device.

#### M4. Illumina barcoded sample preparation

All DNA oligonucleotide preparations (single-stranded DNA synthesised by IDT) were resuspended in nuclease-free water and double-stranded using either `AmpliSeq` or `RevSeq` (with either the RV_20_ or the N_9_ primers), the resulting amplified or reverse-complemented products were cleaned-up with two consecutive steps of 80% EtOH, after mixing the sample with a 180 %(v/v) ratio of AMPure XP beads (Beckman, US). Next, Library preparation was performed using the instructions from NEBNext UltraExpress® DNA Library Prep Kit and barcoding with NEBNext® Multiplex Oligos for Illumina® (96 Unique Dual Index Primer Pairs) in an equivalent manner as with its homologue ONT samples.

#### M5. Strand-aware alignment using anchor-based read filtering for the assessment of oligo quality

The code for strand-aware sequence alignment and logo generation workflow used for the generation of sequence logo frequency plots used in this work for sequence analysis is available in the Supplementary Material SS1 (Script S1).

For strand-aware alignment using anchor-based read filtering after DNA sequencing using an MinION M1kc device (ONT, UK), the sequencing reads, separated by each barcode (SQK-NBDD114.24 - ONT, UK), as `.fastq` files with a Q score equal or greater than 9, were retrieved from the MinION Mk1C (ONT, UK) device. The reads contained within `.fastq` files were then screened against a reference sequence corresponding to either the 75 nt sequence, the 90 nt sequence, or the 150 nt sequence (see `ref_sequence.fa` in the bash command below) using the executed bash command with Script S1:

~~~
python strand_aware_logo.py
-fastq_dir fastq_pass/
--reference ref_sequence
--anchor_len 20
--output logo.png
~~~

Sequencing reads were then processed to generate strand-aware sequence logos relative to the reference sequence to quality check against (see referene sequence in Figure 5). Script S1 gets as input the directory containing `.fastq` files (i.e the totality of sequencing reads), a reference sequence in `.fasta` format, and a user-defined anchor length (20 nt in length in this work) specifying the number of nucleotides at each end of the reference to serve as alignment anchors. The reference sequence was first read from the `.fasta` file, and all sequencing reads were extracted from the provided `.fastq` file-sequencing-containing folder.

To account for strand orientation, each read was compared to the reference sequence in both the forward and reverse complement orientations. Reads were first checked for a match to the left (5’ end) anchor of the reference; if a match was found, the read was aligned accordingly. Reads that did not match the left anchor were reverse complemented and then checked for a match to the right (3’ end) anchor of the reference. Only reads matching either the left or right anchor were retained for sequence logo frequency analysis. These selected reads were then padded or trimmed as necessary to match the length of the reference sequence, ensuring consistent alignment across all positions.

For the aligned reads, a position-by-position count of each nucleotide (`A`, `C`, `G`, `T`, and `-`gaps) was computed without penalisation for any insertions, deletions or substitutions, with the goal to reflect the reality of the oligo sample. These counts were normalised to generate frequency matrices, which were visualised as sequence logos using the *logomaker* Python library[17]. The resulting logo represents nucleotide variability at each position relative to the reference and is proposed as a measure of quality, taking into account each of the methods used (`AmpliSeq`/PCR or `RevSeq`, with or without random primers). The script also identifies and displays the most abundant sequence, the reference sequence, and a consensus sequence (defined as the most frequent nucleotide at each position, or ‘`N`’ if no nucleotide is dominant). The final output was saved as a `.png` image containing these visualizations, and summary of sequences passing the anchoring filters used for the frequency sequence logo plots (Figure 5 are plotted in Figure 3 which include the number of reads processed, the most abundant sequence, and the consensus sequence.

In short, this pipeline (Script S1, Figure 7) processes sequencing reads by matching forward reads to a user-defined left anchor and reverse-complemented reads to a right anchor, both derived from a reference sequence, which can correspond to the primers used for PCR amplification (`AmpliSeq`) or reverse complementation using RV_20_ (`RevSeq`). Afterwards, reads that align (with no penalisation algorithms) successfully are trimmed and aligned based on anchor positions to ensure positional consistency. The processed reads are then used to compute a sequence logo that reflects nucleotide frequency at each position.

#### M6. Second-strand synthesis using the Klenow fragment of the DNA polymerase I

For reverse-complementation (`RevSeq`) using the low-temperature Klenow fragment of the DNA polymerase I, using random nonamers (N_9_) (Figure 2) as priming sequences (`N9-Klen` samples, Figures 3 and 5), a sample of 200 fmol of ssDNA was mixed in 5 *µ*L of 10X rCutsmart Buffer (NEB, UK) 1 *µ*L of a 100 *µ*M solution of random nonamers (N_9_), 2 *µ*L of a 10 mM solution of dNTPs (NEB, UK) and 38.85 *µ*L of nuclease-free water. The mixture was annealed heating the mixture at 94 ºC and down to 25 ºC over a period of 10 minutes. Afterwards, 0.15 *µ*L of a 50 mM solution of NAD+ (NEB, UK) and 2 *µ*L of the Klenow fragment (exo-) of the DNA polymerase I (M0212S - NEB, UK) at 5, 000 U/*µ*L was added to the mixture before incubation at 20 ºC for over an hour.

#### M7. Amplification or second-strand synthesis using HiFi polymerases

For reverse-complementation (`RevSeq`) using RV_20_ primers (Figure 2) and the Q5 DNA polymerase (NEB, UK), a sample of 200 fmol of ssDNA was mixed in 22.5 *µ*L of nuclease-free water, 2.5 *µ*L of a 10 *µ*M reverse-complementary primer (*i*.*e*. referred to as the RV_20_) to the 5-end of the ssDNA sequence, and 25 *µ*L of 2X Q5 Master Mix (NEB, UK). The mixture was incubated following a standard PCR themocycling protocol (as recommended by NEB manufacturing instructions) over 24 cycles. For `AmpliSeq`, a 20.5 *µ*L of nuclease-free water was used to dissolve the 200 fmol ssDNA, and both 2.5 *µ*L of 10 *µ*M of forward (`FW`) and reverse (`RV`) was used instead.

## ACKNOWLEDGEMENTS

The authors are grateful to Patrycja Sokolowska from New England Biolabs (NEB) for her thorough support, correspondence and troubleshooting to define the protocols showcased in this manuscript, using NEB reagents.

## DECLARATION OF CONFLICT OF INTEREST

The authors declare no conflicts of interest.

## AUTHOR CONTRIBUTIONS

R.R.G. and T.H. conceived the project. R.R.G conceptualised, designed and conducted second-strand synthesis and sequencing experiments, wrote the code for strand-aware alignment and anchor filtering for sequence analysis, analysed the results and wrote the manuscript. A.D. prepared samples and data for comparative Illumina sequencing and analysis. All authors revised the manuscript.

## DATA AVAILABILITY

All sequencing raw `.fastq` files are available in the GitHub repository: https://github.com/ImperialCollegeLondon/OligoSeq Ramirez-Garcia, R. *et al*. | OligoSeq: Rapid nanopore-sequencing of single-stranded oligonucleotides bioR*χ*iv | 15

## Supplementary Material

### Supplementary Section SS1: Strand-aware alignment using anchor-based read filtering

**Script S1**. Code for the strand-aware alignment using anchor-based read filtering for the assessment of oligo quality.

~~~
“““
# # #␣This␣code␣does:
# # # # #␣Match␣forward␣reads␣to␣the␣left␣anchor,
# # # # #␣Match␣reverse␣reads␣(revcomp)␣to␣the␣right␣anchor,
# # # # #␣Combine␣them␣into␣one␣sequence␣logo␣plot␣to show␣varaibility
# # # # #␣Usage␣example:
␣␣␣␣␣␣␣␣python␣strand_aware_logo.py␣--fastq_dir␣fastq_pass␣--reference␣reference.fa␣--anchor_len␣20␣--output␣logo.png
# # #␣author:␣Robert␣Ramirez-Garcia
“ “ “
**import** os
**import** argparse
**from** collections **import** Counter
**from** Bio **import** SeqIO
**from** Bio.Seq **import** Seq
**import** pandas **as** pd
**import** matplotlib.pyplot **as** plt
**import** logomaker
**import** matplotlib.patches **as** patches
**def** read_fasta(fasta_file):
    *### Reads a* .*fasta file and returns the sequence*
    **with open**(fasta_file, ‘r’) **as file**:
        seq = []
        **for** line in **file**:
            **if not** line.startswith(‘>‘):
                seq.append(line.strip())
    **return** ‘‘.join(seq)
**def** get_sequences_from_fastq_folder(folder):
    *### Reads all sequences from all* .*fastq files within a folder*
    seqs = []
    **for** fname **in** os.listdir(folder):
        **if** fname.endswith(‘.fastq’) **or** fname.endswith(‘.fq’):
            **for** record **in** SeqIO.parse(os.path.join(folder, fname), “fastq”):
                seqs.append(**str**(record.seq))
    **return** seqs
**def** reverse_complement(seq):
    *### Return the reverse complement of a DNA sequence*
    **return str**(Seq(seq).reverse_complement())
**def** anchor_extract(reference, sequence, anchor_len, mode):
    *### Extracts and aligns sequence to reference based on anchor mode*.
    *### Returns a sequence of reference length, padded with ‘-’ if needed*.
    *### Modes: ‘both’, ‘left’, ‘right’*
    ref_len = **len**(reference)
    ref_left = reference[:anchor_len]
    ref_right = reference[-anchor_len:]
    **if** mode == ‘both’:
        **if** sequence.startswith(ref_left) **and** sequence.endswith(ref_right):
            mid_seq = sequence[anchor_len:**len**(sequence)-anchor_len]
            new_seq = ref_left + mid_seq + ref_right
            **return** new_seq[:ref_len].ljust(ref_len, ‘-’)
    **elif** mode == ‘left’:
        **if** sequence.startswith(ref_left):
            mid_seq = sequence[anchor_len:]
            new_seq = ref_left + mid_seq
            **return** new_seq[:ref_len].ljust(ref_len, ‘-’)
    **elif** mode == ‘right’:
        **if** sequence.endswith(ref_right):
            mid_seq = sequence[:**len**(sequence)-anchor_len]
            new_seq = mid_seq + ref_right
            **return** new_seq[-ref_len:].rjust(ref_len, ‘-’)
    **return** None
**def** get_most_abundant_sequence(seq_list):
    **if not** seq_list:
        **return ““
          return** Counter(seq_list).most_common(1)[0][0]
**def** create_sequence_logo(seq_strings, reference, output_file=“sequence_logo.png”):
    max_length = **len**(reference)
    alphabet = [‘A’, ‘C’, ‘G’, ‘T’, ‘-’]
    counts_matrix = pd.DataFrame(0, index=**range**(max_length), columns=alphabet)
    **for** seq **in** seq_strings:
        **for** pos, char **in enumerate**(seq):
            **if** char **in** alphabet:
                counts_matrix.loc[pos, char] += 1
    freq_matrix = counts_matrix.div(counts_matrix.**sum**(axis=1), axis=0).fillna(0)
    consensus = ‘‘
    **for** pos **in range**(max_length):
        max_freq = freq_matrix.iloc[pos].**max**()
        **if** max_freq > 0.5:
            consensus += freq_matrix.iloc[pos].idxmax()
        **else:**
            consensus += ‘N’
    colour_code = {
        ‘A’: ‘#008000’, # *Green*
        ‘C’: ‘#0000FF’, # *Blue*
        ‘G’: ‘#FFA500’, # *Orange*
        ‘T’: ‘#FF0000’, # *Red*
        ‘-’: ‘#808080’, # *Gray*
        ‘N’: ‘#808080’
    }
    most_abundant = get_most_abundant_sequence(seq_strings)
    fig, (ax_logo, ax_abundant, ax_ref, ax_cons) = plt.subplots(
        4, 1, figsize=(15, 10),
        gridspec_kw={‘height_ratios’: [4, 1, 1, 1]},
        sharex=True
    )
    logo = logomaker.Logo(freq_matrix, ax=ax_logo, color_scheme=colour_code)
    logo.style_spines(visible=False)
    ax_logo.set_ylabel(‘Frequency’)
    ax_logo.set_title(‘Sequence ␣Logo’)
    **for** pos, char **in enumerate**(most_abundant):
        **if** char **in** colour_code:
            colour = colour_code[char]
            rect = patches.Rectangle((pos-0.5, 0), 1, 1, facecolor=colour, alpha=0.6)
            ax_abundant.add_patch(rect)
            ax_abundant.text(pos, 0.5, char, ha=‘center’, va=‘center’, fontsize=10, fontweight=‘bold’, color=‘white’)
    ax_abundant.set_ylim(0, 1)
    ax_abundant.set_yticks([])
    ax_abundant.set_title(‘Most ␣Abundant␣Sequence’)
    ax_abundant.set_frame_on(False)
    **for** pos, char **in enumerate**(reference.ljust(max_length, ‘-’)):
        **if** char **in** colour_code:
            colour = colour_code[char]
            rect = patches.Rectangle((pos-0.5, 0), 1, 1, facecolor=colour, alpha=0.6)
            ax_ref.add_patch(rect)
            ax_ref.text(pos, 0.5, char, ha=‘center’, va=‘center’, fontsize=10, fontweight=‘bold’, color=‘white’)
    ax_ref.set_ylim(0, 1)
    ax_ref.set_yticks([])
    ax_ref.set_title(‘Reference ␣Sequence’)
    ax_ref.set_frame_on(False)
    **for** pos, char **in enumerate**(consensus):
    **if** char **in** colour_code:
        colour = colour_code[char]
        rect = patches.Rectangle((pos-0.5, 0), 1, 1, facecolor=colour, alpha=0.6)
        ax_cons.add_patch(rect)
        ax_cons.text(pos, 0.5, char, ha=‘center’, va=‘center’, fontsize=10, fontweight=‘bold’, color=‘white’)
    ax_cons.set_ylim(0, 1)
    ax_cons.set_yticks([])
    ax_cons.set_xlabel(‘Position’)
    ax_cons.set_title(‘Consensus ␣Sequence’)
    ax_cons.set_frame_on(False)
    plt.tight_layout()
    plt.savefig(output_file, dpi=300)
    **print**(f”Most␣abundant␣sequence: ␣{most_abundant}”)
    **print**(f”Consensus␣sequence: ␣{consensus}”)
    **print**(f”Sequence␣logo␣saved␣as␣{output_file}”)
  **if** __name__ == “__main__”:
    parser = argparse.ArgumentParser(description=“Strand-aware␣anchor␣alignment␣and␣sequence␣logo␣generator.”)
    parser.add_argument(“--fastq_dir”, **type=str**, default=“fastq_pass”, **help**=“Directory␣ ␣fastq␣files”)
    parser.add_argument(“--reference”, **type=str**, required=True, **help**=“.fasta␣/ ␣.fa␣file␣with ␣reference␣sequence”)
    parser.add_argument(“--anchor_len”, **type=int**, default=20, **help**=“Length␣of␣anchor␣region␣ (nt)”)
    parser.add_argument(“--output”, **type=str**, default=“sequence_logo.png”, **help**=“Output␣image␣filename”)
    args = parser.parse_args()
    reference = read_fasta(args.reference)
    all_seqs = get_sequences_from_fastq_folder(args.fastq_dir)
    **print**(f”Total␣sequences␣read: ␣{len(all_seqs)}”)
    selected = []
    **for** seq **in** all_seqs:
        aligned = anchor_extract(reference, seq, args.anchor_len, mode=“left”)
        **if** aligned:
            selected.append(aligned)
            **continue**
        rev_seq = reverse_complement(seq)
        aligned_rc = anchor_extract(reference, rev_seq, args.anchor_len, mode=“right”)
        **if** aligned_rc:
            selected.append(aligned_rc)
    **print**(f”Sequences␣passing␣anchor␣filter: ␣{len(selected)}”)
    **if not** selected:
        **print**(“No␣sequences␣passed␣the␣anchor␣filter.”)
        exit(1)
    *### Precomputes most abundant and consensus for clean reporting*
    most_abundant = get_most_abundant_sequence(selected)
    max_length = **len**(reference)
    counts_matrix = pd.DataFrame(0, index=**range**(max_length), columns=[‘A’, ‘C’, ‘G’, ‘T’, ‘-’])
    **for** seq **in** selected:
        **for** pos, char **in enumerate**(seq):
            **if** char **in** counts_matrix.columns:
                counts_matrix.loc[pos, char] += 1
    freq_matrix = counts_matrix.div(counts_matrix.**sum**(axis=1), axis=0).fillna(0)
    consensus = ‘‘
    **for** pos **in range**(max_length):
        max_freq = freq_matrix.iloc[pos].**max**()
        **if** max_freq > 0.5:
            consensus += freq_matrix.iloc[pos].idxmax()
        **else:**
            consensus += ‘N’
    *### Outputs all expected logs*
    **print**(f”Most␣abundant␣sequence: ␣{most_abundant}”)
    **print**(f”Reference␣sequence␣used␣in␣plot: ␣{reference}”)
    **print**(f”Consensus␣sequence: ␣{consensus}”)
    *### Creates and saves the logo*
    create_sequence_logo(selected, reference, args.output)
~~~

### Supplementary Section SS2: Sequencing costs per platform

#### SS1. ONT sequencing costs

For example, running a sample through using ONT requires the purchase of flow cells or flongles (in this study, R10 flongles from ONT were used). The cost of each flongle is US$67.5 while the native barcoding kit (SQK-NBD114.24 Native Barcoding Kit 24 V14) used in this work costs about US$699.00 for a set of 6 reactions with each reaction able to barcode up to 24 different sequences, which falls to US$4.85 per barcode. Thus the total cost of sequencing one single oligo falls to US$7.7 per oligo (provided a full set of 24 oligoes are sequenced per flongle). To this is important to add the cost of reagents to convert the single-stranded oligonucleotides to double-stranded which costs US$1.28 per reaction (£0.95 according to current NEB UK prices for the reagents involved making a maximum of US$9 per oligo sequenced, calculated as follows:

- ONT R10 Flongle: US$67.50 or £51.00 (per flow cell), amortized to US$2.81 or £2.13 per sample when 24 samples are multiplexed
- ONT Native Barcoding Kit (SQK-NBD114.24): US$699.00 or £529.00 for 6 reactions (24 barcodes), equating to US$4.85 or £3.67 per sample
- Double-stranding reagents (NEB): US$1.28 or £0.95 per reaction

Total cost per oligo: US$8.94 or £6.75, when sequencing 24 samples per Flongle, considering an exchange rate of 1.33 US$/£ (as of December 2025).

### SS2. PacBio sequencing costs

The SMRTbell® adapter index plate 96A costs £571 for 96 samples, equating to £5.95 per sample. The SMRTbell® Prep Kit 3.0 (PacBio® 102-182-700), used to circularize DNA, costs £1, 598 and supports 24 libraries of 96 samples each, enabling up to 2,304 samples with a per-sample cost of £0.69 if fully utilized. The Binding Kit 3.1 and Cleanup Beads 3.1 (PacBio® 102-194-200), required for polymerase binding, is priced at £1, 434, also supporting 2,304 samples and reducing the per-sample cost to £0.62 under full usage. Additional sequencing costs include a flow cell at £3, 584 for four runs (£896 per run), translating to £9.33 per sample; SMRTcell Oil 4x (5 tubes for 4 runs) costs £158, adding £0.411 per sample; lids for oil tubes cost £33 for 20 runs, adding £0.086 per sample; and the Sequel II Sequencing Kit 2.0 for four flow cells costs £761, adding £1.98 per run. Additional overheads include costs for specific plates, tips, and lids required for PacBio equipment operation. Summing all up, the total cost per sample, excluding final overheads (plates, tips, lids), is £18.45, calculated as follows:

- SMRTbell^®^ adapter index plate 96A: £5.95 or US$7.91
- SMRTbell^®^ Prep Kit 3.0: £0.69 or US$0.92
- Binding and Cleanup Kit 3.1: £0.62 or US$0.82
- Flow cell: £9.33 or US$12.41
- SMRTcell Oil: £0.411 or US$0.55
- Oil lid: £0.086 or US$0.11
- Sequel II Sequencing Kit 2.0: £1.98 or US$2.63

Which, based on a £-to-USD exchange rate of 1.33 (December 2025), costs an average (much) greater than £18.45 or US$24.55 per sample to use PacBio (in the unlikely event that over 2,000 samples are ready to be processed).

#### SS3. Illumina sequencing costs

This workflow for Illumina sequencing is similar to the one described for ONT, which consists in (1) adapter ligation and (2) barcoding ligation, and also needs a proprietary flow cell for use. This workflow also needs double-stranding of single-stranded oligonucleotides which includes the overhead of US$1.28 per reaction/sample (£0.95 according to current NEB UK prices for the reagents involved, For sample specific-preparation for IllUmina sequencing, the NEBNext UltraExpress^®^ DNA Library Prep Kit is used, that includes dA tailing and end-repair, adapter ligation, PCR enrichment and cleanup, the cost for a kit capable of processing 96 samples is £1, 880.00 which falls down to £19.58 per sample. For multiplexing 96 reactions for a separate sample per barcode, it is possible to use (similarly as with the ONT native barcoding kit) the barcoding kit corresponding to Illumina (NEBNext® Multiplex Oligos for Illumina® (96 Unique Dual Index Primer Pairs)) where a whole plate is £570.00 which would be £5.93 per sample. Once the sample is prepared using these reagents, a flow cell is needed for sequencing which costs no less than USD129.00 and using an external serivce provider Novogene (Cambridge, United Kingdom) costed £111.36.

- i1 Flow Cell: US$129.00 or £96.99 (per flow cell), amortized to US$1.78 or £1.34 per sample when 96 samples are multiplexed
- NEBNext UltraExpress^®^ DNA Library Prep Kit: US$2, 500.40 or £1, 880.00 for 96 reactions, equating to US$26.04 or £19.58 per sample
- NEBNext^®^ Multiplex Oligos for Illumina^®^ (96 Unique Dual Index Primer Pairs): US$758.10 or £570.00 for 96 reactions, equating to US$7.89 or £5.93 per sample
- Double-stranding reagents (NEB): US$1.28 or £0.95 per reaction

Which, based on a £-to-US$ exchange rate of 1.33 (December 2025), costs an average (much) greater than £ $27.80 or US$36.99 per sample to use Illumina iSeq.

#### SS4. HPLC/MS sequencing costs

Conversely, using an Agilent Suite HPLC/MS service at Imperial College London which runs oligonucleotides on the Oligo LC/MS - 1290 Infinity III with Pro iQ Plus SQ MS (Agilent Technologies, California, USA), costs no less than US$43 per hour and at least (with 0.5 hours dedicated to initial equipment start-up, heat up and sample preparation and 0.33 hours per oligo processed) will be required to run 3 different oligo samples. Without considering the reagents needed for operation and maintenance. Thus the cost of running one oligo falls to at least US$21.5 + US$14.33 = US$36 per oligo. Still, without considering equipment start-up timings, the cost of adding up one more oligo in the HPLC/MS analysis is US$14.33, which is a 100 % greater than using ONT technology, and the HPLC/MS in question is only able to resolve molecules by molecular mass which adds further overhead complexity to resolve the actual DNA sequences, increasing its cost therefrom. Thus the only foreseeable advantage could be the lack of sample preparation—as the ssDNA oligo samples can be run directly on the HPLC/MS equipment—which might help save time, however the need to run the mobile phases and heat the colum for HPLC/MS overrides this seeming advantage of a lack of sample-prep, thus the summary costs follow below as:

- HPLC/MS instrument time (start-up): US$21.50 or £16.10 (0.5 hours at US$43/hour)
- HPLC/MS run per oligo: US$14.33 or £10.73 (0.33 hours per oligo at US$43/hour)

Total cost per oligo: US$35.83 or £26.83, excluding reagents and maintenance considering an exchange rate: 1.33 US$/£.

### Supplementary Section SS3: Supplementary figures

**Fig. S1.**
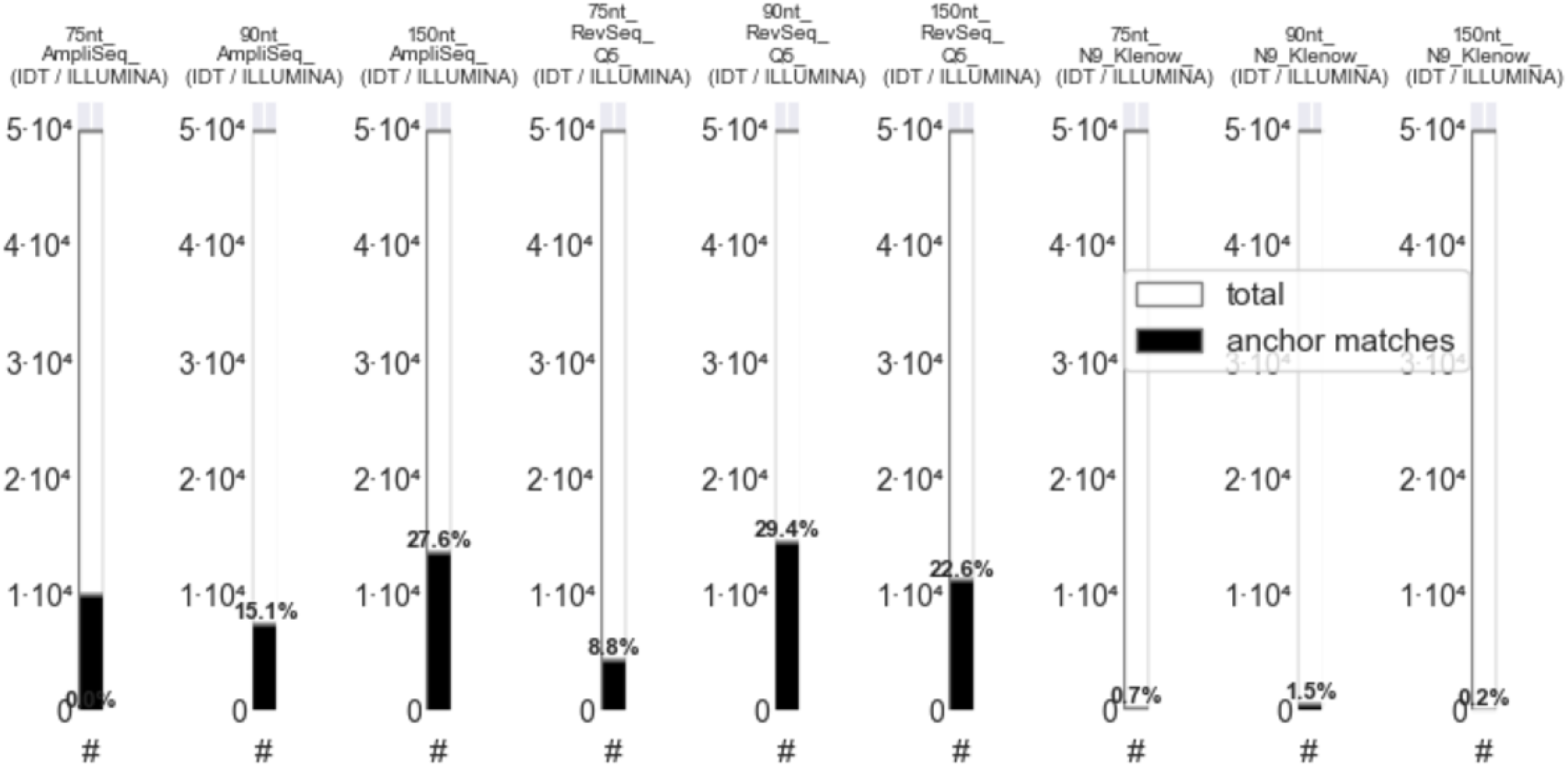
Number (#) of sequence anchor matches that can be aligned to target sequence with Illumina sequencing. **Total:** Total number of reads extracted from the sequencing run. **Anchor matches:** Sequences that could be aligned to a reference sequence through anchoring reads to the first 5’ end 20 nt of the plus `(+)` and minus `(−)` DNA sequences filtered using Script S1, regardless of % identity beyond the 20 nt anchors. All experiments were normalised to 200 fmol per load of dsDNA (`AmpliSeq` and `RevSeq`, either with Q5 or N9-Klen) or ssDNA (Control). Substrate ssDNA for all samples was synthesised using an *in-house* Kilobaser or IDT before double-stranding using enzymes or annealing. For a comparison of the results here obtained with those with various reads using ONT sequencing see Figure 3.

**Fig. S2.**
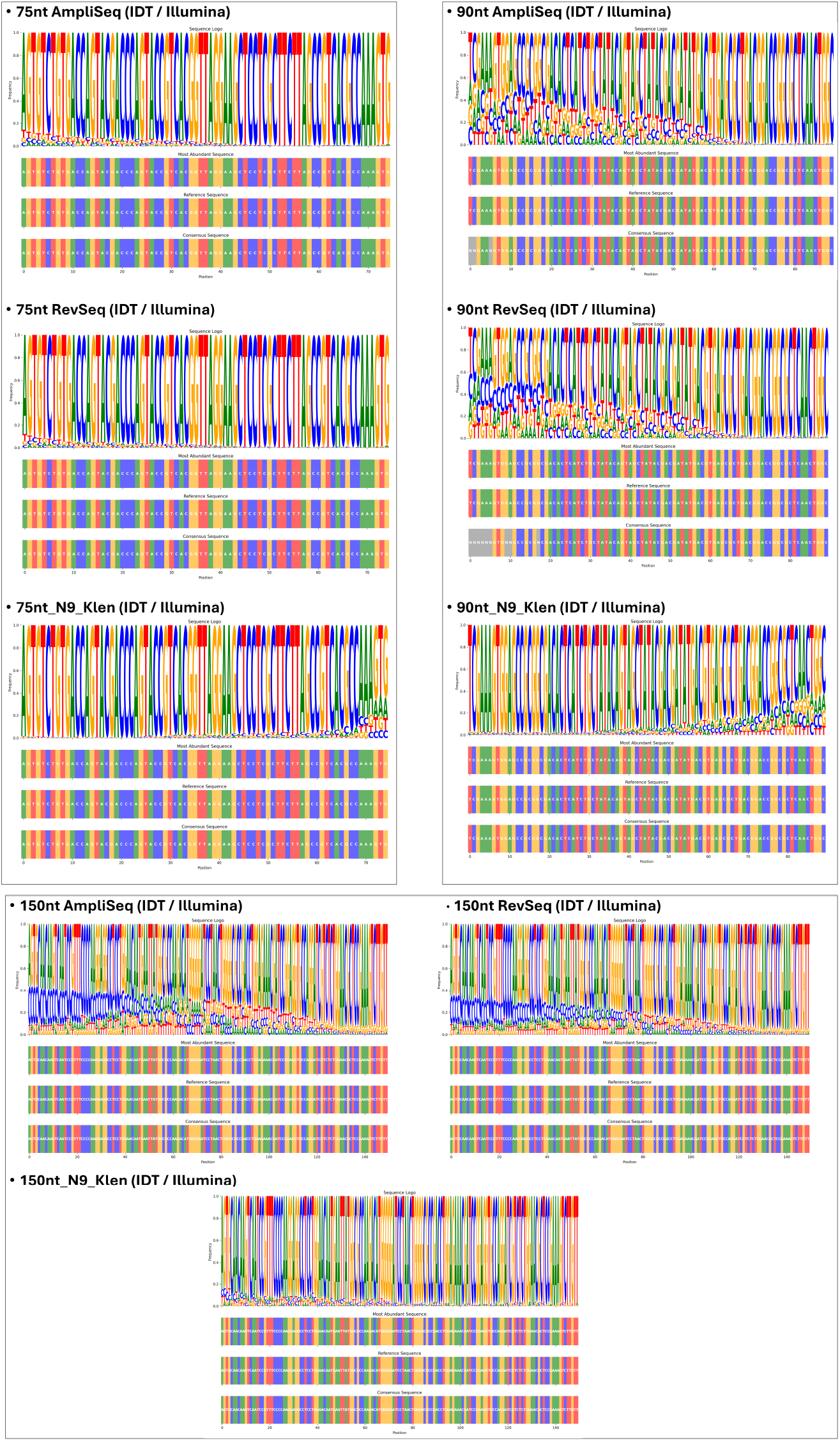
Sequence logo plots for the assessment of the oligo quality of the equivalent 75-mer, 90-mer and 150-mer samples, sequenced using Illumina instead of ONT.

## Notes

https://github.com/ImperialCollegeLondon/OligoSeq

### Competing Interest Statement

The authors have declared no competing interest.

### Summary of Updates

This revision now includes comparative experimental data includes a set of experiments with Illumina sequencing using the same samples so that the readers can have a vision of the results analysed in terms of anchor read matches and sequence variability, now using Illumina to compare results with results with ONT/nanopore sequencing technology. The new version of this manuscript also includes now a polishing of the text and has deleted paragraphs with redundant information. Additionally, plots have now been separated into several figures in order to facilitate the visualisation.

